# Salinomycin inhibits epigenetic modulator EZH2 to enhance Death Receptors in Colon Cancer Stem Cells

**DOI:** 10.1101/2020.02.03.932269

**Authors:** Anup Kumar Singh, Ayushi Verma, Akhilesh Singh, Rakesh Kumar Arya, Shrankhla Maheshwari, Priyank Chaturvedi, Mushtaq Ahmad Nengroo, Krishan Kumar Saini, Achchhe Lal Vishwakarma, Kavita Singh, Jayanta Sarkar, Dipak Datta

## Abstract

Drug resistance is one of the trademark features of Cancer Stem Cells (CSCs). We and others have recently shown that paucity of functional death receptors (DR4/5) on the cell surface of tumor cells is one of the major reasons for drug resistance, but their involvement in the context of in CSCs is poorly understood. By harnessing CSC specific cytotoxic function of salinomycin, we discovered a critical role of epigenetic modulator EZH2 in regulating the expression of DRs in colon CSCs. Our unbiased proteome profiler array approach followed by ChIP analysis of salinomycin treated cells indicated that the expression of DRs, especially DR4 is epigenetically repressed in colon CSCs. Concurrently, EZH2 knockdown demonstrated increased expression of DR4/DR5, significant reduction of CSC phenotype such as spheroid formation *in-vitro* and tumorigenic potential *in-vivo* in colon cancer. TCGA data analysis of human colon cancer clinical samples shows strong inverse correlation between EZH2 and DR4. Taken together, this study provides an insight about epigenetic regulation of DR4 in colon CSCs and advocates that drug resistant colon cancer can be therapeutically targeted by combining TRAIL and small molecule EZH2 inhibitors.

## 1. Introduction

According to global cancer statistics data, colorectal cancer (CRC) is the fourth most common malignancy worldwide, with 1.09 million new incidences diagnosed in 2018^1^. The standard care for CRC patients is surgical resection followed by adjuvant therapy with chemotherapeutics or molecular targeted therapy. The therapy failure in terms of disease relapse and metastasis often results from the presence of drug-refractory cell population, present in highly heterogeneous solid tumors^2, 3^. Increased epithelial-mesenchymal-transition (EMT), stemness and cellular-plasticity are major contributory factors for the development apoptosis resistance in cancer cells. Recently, in two different contexts, we have shown that inducing lethal autophagy is an effective strategy to overcome apoptosis resistance in colon cancer^4^ and SPINK-1 regulates cellular plasticity and stemness in prostate cancer^5^. The phenotype of drug resistance cells in solid tumors like colon cancer is extensively overlapping with the properties of cancer like stem cells or cancer stem cells (CSCs) that demonstrate enormous cellular plasticity, self-renewal and tumor-initiating capabilities^6^. Gupta et al., in their pioneering work, first identified salinomycin as a potent CSC killer by utilizing high throughput screening approach^7^. Due to its immense clinical potential, this finding itself fueled exponential growth in the literature of salinomycin and CSCs; and some of them proposed its mode of actions in different cancers via alteration of different signaling cascades^8^. Numerous recent studies emphasize the crucial role of epigenetic regulation in modulating various CSC attributes including drug resistance^9–11^. Therefore, how salinomycin is able to epigenetically target them is still remained elusive. Histone modifications through Polycomb Group (PcG) of proteins have shown to drive stem cell biology via chromatin remodeling, specifically by catalyzing posttranslational modifications on histone proteins via Polycomb Repressive Complex 1 (PRC1) and PRC2^12–14^. Enhancer of zeste homologue 2 (EZH2) is the catalytic component of the PRC2 complex that is involved in transcriptional repression of its target gene by tri-methylation of lysine 27 at histone H3 (H3K27)^15^. Seminal studies by Chinnaiyan’s group first linked EZH2 with the progression of solid tumors^16, 17^. More recently, EHZ2 has shown to regulate hallmarks of CSC properties like self-renewal, tumorigenic potential and drug resistance in different tumor types^18, 19^. Most important breakthrough is EZH2 inhibitor Tazemetostat (EPZ6438) just received FDA approval for metastatic epithelioid sarcomas.

We and others have shown that paucity of functional DR4/DR5 on the cell surface of colon tumor cells is one of the major reasons for drug resistance in general, but their involvement in the context of drug resistance in CSCs are not known so far^20–23^. By harnessing CSC specific cytotoxic function of salinomycin, we discovered an unprecedented role of EZH2 in modulating the expression of death receptor in colon cancer. Our unbiased proteome profiler array approach indicated salinomycin mediated regulation of death receptors in colon cancer. Further molecular insight of death receptor regulation by salinomycin suggests that expression of DR4, DR5 was found to be epigenetically repressed in colon CSCs due to the EZH2 mediated histone hyper-methylation, independent of promoter DNA methylation. Salinomycin or EZH2 inhibitor treatment withdraws H3K27 trimethylation marks from the promoter of death receptors and up-regulates the functional expression of DR4 and DR5 and sensitizes colon cancer cells against TRAIL therapy. Concurrently, in a similar setting, genetic loss of EZH2 resulted in significant reduction in CSC properties *in-vitro* and *in-vivo*. Taken together, this study provides a novel link for epigenetic regulation of death receptor in colon CSCs that can be therapeutically targeted in future.

## 2. Materials and Methods

### 2.1 Reagents and antibodies

Salinomycin, Cisplatin, GSK-343, DZNep, 5-AZA, TSA, Dimethyl Sulfoxide (DMSO), Sulforhodamine B (SRB) and Bovine Serum Albumin (BSA) were purchased from Sigma-Aldrich. EPZ6438 was purchased from Apex biosciences. Puromycin was purchased from thermo fisher scientific. Cisplatin (1mg/ml) solution brought from CADILA. APC and PE conjugated CD133 (clone 1); and PE conjugated DR4 and DR5 antibodies were obtained from Miltenyi Biotec and E-Biosciences respectively. Antibodies for DR5, SP1, GAPDH, p53, YY1, β-actin, DNMT1, EZH2, HDAC1, HDAC6, SUZ12, RING1A, BMI1, H3K27me3, used in the western blotting were purchased from Cell Signaling Technology. Anti DR4 antibody, PVDF membrane and stripping buffer, were procured from Millipore Inc. Aldefluor kit was bought from Stem cell technologies. Magnetic ChIP kit (Cell Signaling Technology), Verso one step RT-PCR kit, BCA protein estimation kit, RIPA cell lysis buffer, blocking buffer, Super Signal West Pico and Femto chemiluminescent substrate, Alexafluor 488 conjugated Annexin-V, Pure Link^TM^ RNA Mini kit, Lipofectamine-2000, Alexafluor 488/594 conjugated secondary antibodies, FBS, DMEM, RPMI-1640 media, Anti-Anti, were purchased from Thermo Fisher Scientific. Proteome profiler Human Apoptosis array kit and recombinant human TRAIL were obtained from R&D Systems. Primers for *GAPDH, DR4, DR5* genes and those used in ChIP assay were purchased from IDT Inc. All chemicals were obtained from Sigma unless specified otherwise.

### 2.2. Cell culture and treatment

Representative colon cancer cell lines DLD-1, RRID: CVCL_0248 (epithelial, non-metastatic colorectal adenocarcinoma),SW-620, RRID:CVCL_0547 (Dukes’ type C, metastatic colorectal adenocarcinoma) and LOVO, RRID:CVCL_0399 (grade IV, colorectal adenocarcinoma), HT-29, RRID:CVCL_0320 (colorectal adenocarcinoma) were obtained from American Type Culture Collection (ATCC), USA. Mycoplasma free early passage cells were resuscitated from liquid nitrogen vapor stocks and inspected microscopically for stable phenotype before use. All cell lines used in the study are authenticated by STR profiling. Cells were cultured as monolayers in recommended media supplemented with 10% FBS, 1-X anti-anti (containing 100 μg/ml streptomycin, 100 unit/ml penicillin and 0.25 μg/ml amphotericin B) and maintained in 5% CO_2_ and humidified environmental 37°C. In all treatments, salinomycin and GSK-343 and EPZ6438were dissolved in cell culture grade DMSO at concentration of 10 mM and Cisplatin 1mg/ml. TSA, 5’-Azacytidine, DZNep, TRAIL and other chemicals were dissolved as per supplier’s recommendation. The sub-confluent cells were treated with required doses of compounds in all the experiments.

### 2.3 Cell Viability Assay

A standard colorimetric SRB assay was used for the measurement of cell cytotoxicity^24^. In this experiment, 10,000 cells of each cell lines were seeded to each well of 96-well plate in 5% serum containing growth medium and incubated overnight to allow for cell attachment. Cells were then treated with test molecule at the required dose and untreated cells received the same volume of vehicle containing medium served as control. After 48 h of incubation, cells were fixed with ice-cold 10% TCA, stained with 0.04% (w/v) SRB in 1% acetic acid, washed and air dried. Bound dye was dissolved in 10mM Tris base and absorbance was measured at 510 nm on a plate reader (Epoch Microplate Reader, Biotek, USA). The cytotoxic effects of compounds were calculated as % inhibition in cell growth as per the formula [100-(Absorbance of compound treated cells/ Absorbance of untreated cells)] X 100.

### 2.4 Flowcytometry staining and analysis

For Annexin-V and CD133 double staining, cells were trypsinized and washed with 1× binding buffer and then incubated with FITC conjugated Annexin-V (BD Biosciences) for 15 min at room temperature in the dark. After Annexin-V staining, cells were washed with PBS containing 0.1 % FBS, and then incubated with either appropriate isotype control antibody or PE conjugated CD133/1 on ice in the dark for 15 min. After washing with PBS, cells were acquired in FACS-Calibur^TM^ (BD Biosciences) and analyzed by using FlowJo software (Tree Star Inc, USA). For DR4/DR5 and CD133 dual staining, we used PE conjugated DR4/DR5 and APC conjugated CD133 antibodies along with isotype controls and followed above mentioned protocol for FACS analysis. The ALDEFLUOR kit (StemCell Technologies, USA) was used to assess the population with a high ALDH enzymatic activity. Cells were suspended in ALDEFLUOR assay buffer containing ALDH substrate (BAAA, 1 μmol/l per 1×10^6^ cells) and incubated during 40 min at 37°C. For each sample of cells, an aliquot was treated with 50mmol/L diethylaminobenzaldehyde (DEAB), a specific ALDH inhibitor as negative control. Finally, cells were acquired and analyzed by following the same protocol.

### 2.5 Human apoptosis protein array

Apoptosis array analysis was performed using the Proteome Profiler Human Apoptosis Array Kit (ARY009) from R&D Systems according to the manufacturer’s instructions. Briefly, SW620 cells were plated at a density of 1× 10^6^ cells in a 60 mm tissue culture dish for 24 hours and then were treated with 10μM concentration of salinomycin. After 24 hours of treatment, cell lysates were prepared by homogenization in lysis buffer 16 (R&D systems) and incubated on ice for 30 min. Lysates were cleared by centrifugation at 14,000 g for 15 min at 4 °C. The protein concentration of the supernatants was measured using the BCA protein assay kit (Pierce) with bovine serum albumin as the standard. Apoptosis array was performed as described previously^25^.

### 2.6 RNA isolation and Reverse Transcription-PCR (RT-PCR) analysis

Total RNA was prepared using the Pure Link ^TM^ RNA Mini kit The cDNA synthesis and PCR were carried out by Verso one-step reverse transcription-PCR (RT-PCR) kit using gene-specific primers and following the manufacturer’s protocol, and the detailed procedure as described previously^26^. The sequence of the oligonucleotide primers used are as follows: for human GAPDH: Forward- 5’-GTCAGTGGTGGACCTGACCT-3’, Reverse- 5’-AGGGGAGATTCAGTGTGGTG-3; product size-395bp, for human DR4: Forward-5’-AGAGAGAAGTCCCTGCACCA-3’,Reverse5’AGAGAGAAGTCCCTGCACCA-3’; product size-366bp, for human DR5: Forward-5’-TGCAGCCGTAGTCTTGATTG-3’,Reverse5’GCACCAAGTCTGCAAAGTCA-3’; product size-389bp. TaqMan gene expression assay from Thermo Fisher scientific were used for DR4(Assay ID: <u>Hs00269492_m1</u> DR5 (Assay ID: <u>Hs00366278_m1</u>) gene amplification.

#### Chromatin Immune-Precipitation (ChIP) Assay

ChIP assay was conducted using the Simple ChIP(R) Enzymatic Chromatin IP Kit purchased from Cell Signaling Technology following the manufacturer’s instruction. In brief, nearly half confluent SW620 cells were treated with 10 µM salinomycin, EPZ643810μM. After 24 hours of treatment 10 millions of cells were taken from each group, genomic DNA and protein were cross-linked by addition of formaldehyde (1% final concentration) directly into the culture medium and incubated for 10 min at 37°C. Cells were then collected and lysed in 200 μl of membrane extraction buffer containing protease inhibitor cocktail followed by 20U of micrococcus nuclease (MNase) treatment in digestion buffer to obtain chromatin fragments. Cells were sonicated for 7 minutes at the 30% of amplitude with a cycle of 30 second on followed by 30 second off to generate DNA fragments of 200 to 500 bps long. After centrifugation, the cleared supernatant was diluted 10-fold with IP buffer and 10 µl of it was kept as input control and rest is incubated at 4°C overnight with Histone3as positive control, anti-H3K27me3 monoclonal antibody as test for different groups and mouse IgG isotype antibody as negative control. Immune complexes were precipitated, washed, and eluted as per recommended protocol. Finally, DNA-protein cross-linkages were reversed by heating at 65°C for 4 hours; DNA was extracted in phenol/chloroform, precipitated with ethanol, and re-suspended in 50 µl of Tris-EDTA elution of buffer (pH 8.0). Real-time q-PCRs were performed using 2X dyNamo SYBER Green. Primers were designed using Gene Script PCR design software and synthesized by IDT Inc. The sequence of the various primer sets used in the study is as given in the Supplementary Table-1.

### 2.7 Confocal Microscopy

Control and treated cells were fixed with ice-cold pure methanol for 10 min at −20° C followed by blocking with 2% BSA for 1 hour at RT. After overnight primary antibodies (anti-EZH2 and anti-DR5) incubation, cells were washed twice with PBS and incubated with fluorescent-conjugated secondary antibodies at RT for 1 hour, followed by DAPI staining for 5 min at RT. After washing, cells were mounted with anti-fade mounting medium on glass slides and viewed under an inverted confocal laser scanning microscope (Zeiss Meta 510 LSM; Carl Zeiss, Jena, Germany). Plan Apochromat 63X/1.4NA Oil DIC objective lens was used for imaging and data collection. Appropriate excitation lines, excitation and emission filters were used for imaging.

### 2.8 EZH2 knock down by utilizing retroviral transduction

Control and stable EZH2 cell lines were generated by utilizing retroviral mediated transduction system followed by puromycin selection. Control and EZH2 (cat #24230) knockdown retroviral plasmid were brought from Addgene USA. The Phoenix cell line was used for the generation of retroviral particles using the transfection reagent Lipofectamine 2000.The Phoenix cells were plated in the 6-well plate at 80% confluency. Polybrene (8μg/ml) was added to the viral soup during the transduction matured viral particles into the target cells. Cells were subjected to puromycin selection, and the EZH2 knockdown was confirmed by western blot.

### 2.9 *In-vivo* studies in xenograft tumor models

All animal studies were conducted by following standard principles and procedures approved by the Institutional Animal Ethics Committee (IAEC) CSIR-Central Drug Research Institute. Xenograft implantation of colon tumor cells into mice was performed as described earlier^27^. In brief, established stable 1×10^6^ DLD-1 control or EZH2 KD cell in 100 μl PBS were subcutaneously inoculated in to both flanks of the left and right hind leg respectively of each S4-6-week-old nude Crl: CD1-Foxn1^nu^ mice. Throughout the study, tumors were measured with an electronic digital caliper at regular interval and the tumor volume was calculated using standard formula *V* = Π / *6* × *a*^2^ × *b* (*a* is the short and *b* is the long tumor axis). At the end of experiment, mice were sacrificed, and subcutaneous tumors were dissected for further studies.

### 2.10 Analysis of TCGA COAD-colon cancer dataset

IlluminaHiSeq generated transcript data of EZH2 and DR4 of colon cancer patients from the cohort of TCGA-COAD is downloaded from The Cancer Genomic Atlas (TCGA) database by using https://xena.ucsc.edu/ browser. Out of 551 patient samples, only 331 patient samples were showed expression for EZH2 and DR4. Since, EZH2 is over expressed in∼22-25 % of the colon cancer patients in this particular cohort. Eventually, we converted the FPKM values of both EZH2 and DR4 in the form of Log2 FC algorithm. We performed quartile based normalization to segregate the patients based on high and low EZH2 expression^28^. Accordingly, patients corresponding in the top quartile (n=73) (log2 FC >2.5) were considered as EZH2-high whereas, patients in the lower quartile (n=27) (log2 FC< 1.75) were assigned as EZH2-low. The heat map was formed between EZH2 Log2FC values and DR4 Log2FC values by using the heatmapper online software. The Average Linkage was considered as a clustered method and the Euclidean method was considered for distance measurement.

### 2.10 Statistics

Most of the in vitro experiments are representative of at least three independent experiments. Student’s t-test and two-tailed distributions were used to calculate the statistical significance of *in vitro* and *in vivo* experiments. These analyses were done with GraphPad Prism. Results were considered statistically significant when p-values ≤ 0.05 between groups.

## 3. Results

### 3.1 Salinomycin but not cisplatin attenuate cancer stem cell properties and promote apoptosis in colon CSCs

Several recent reports suggested that salinomycin can target CSCs^7, 29^; but whether its cytotoxic effects are selective to CSCs is not well understood. To study the effect of salinomycin on the CSC population in the colon cancer cells, we have selected colon-specific CSC marker (CD133) and Aldehyde dehydrogenase (ALDH) activity as read out for stemness. We checked the expression of CD133 and ALDH activity in different colon cancer cell lines through flow cytometry (Figure 1A and Supplementary Figure 1). The FACS histogram overlays suggest that CD133 expression was least in DLD-1 cells whereas, LOVO, SW620 and HT-29 cells show moderate to high stemness properties (Figure 1A and Supplementary Figure 1). To test the effect of salinomycin on particularly CD133+ versus CD133-cells, we try to first adapt FACS Aria based CD133 sorting to isolate both positive and negative cells from DLD-1 colon cancer cell lines and cultured for couple of days and check the CSC integrity of the cells before salinomycin treatment. Unfortunately, we observed that majority of sorted CD133^+^ cells rapidly converted into CD133^-^ cells and quickly lost their CSC integrity (Supplementary Figure 2), which actually supports the concept of CSC plasticity reported by Chaffer *et.al*, in her classic paper^30^. As CSC sorting in colon cancer cell lines did not offer us right kind of model system, we focus our study on CSC enriched colon cancer cells. Since SW620 cells were showing higher CSC activity, we selected this cell line to study the effect of Salinomycin treatment on CD133 and ALDH activity. Here, we observed that salinomycin significantly reduced CD133 expression as well as ALDH activity in SW620 cells. In contrast, common chemotherapeutic drug cisplatin failed to do so (Figure 1B and 1C).

**Figure 1.**
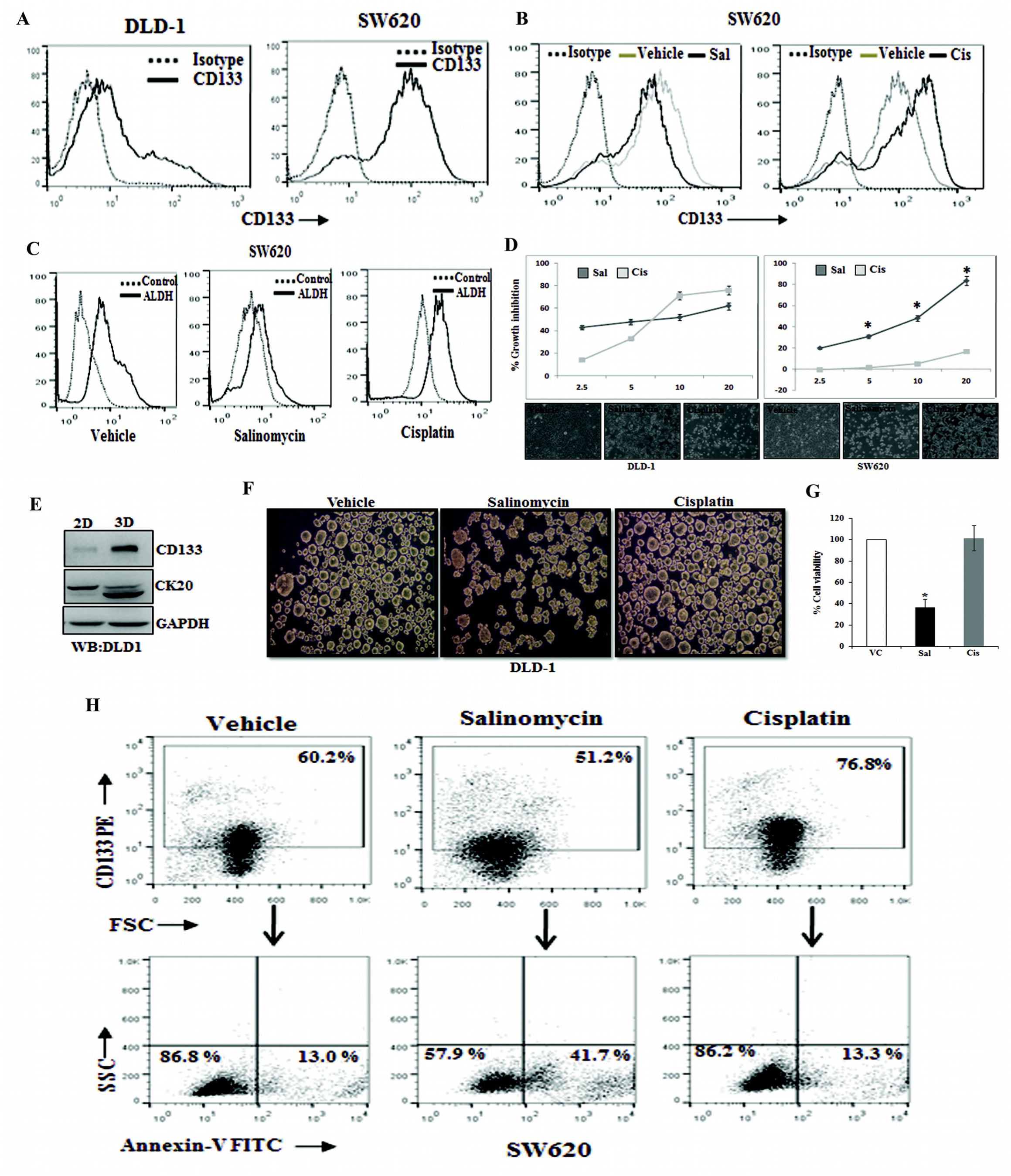
Salinomycin inhibits stemness, reduces spheroids formation and induces apoptosis in colon cancer stem cells. (A) CD133 surface expression was analyzed by FACS in DLD-1, SW620 cells. SW620 cells were treated with either salinomycin (10 µM) or cisplatin (10 µM) for 24 hours and analyzed by FACS for (B) CD133 expression and (C) ALDH activity. (D) DLD-1cells and SW620 cells were treated with multiple doses of salinomycin and cisplatin for 48 hours and subjected to SRB assay to assess their growth inhibitory response. Their corresponding photo micrographs are shown in bottom panel. Data points are average of triplicate readings of samples; error bars, ± S.D. *p < 0.01, compared to cisplatin treated cells. (E) The expression of CD133 and cytokeratin 20 (CK20) in DLD-1 cells were determined in both adherent (2D) and non-adherent (3D) culture by Western blot analysis. (F) DLD-1 cells were plated at 50,000 cells per 9.04 cm^2^ dish in a non-adherent plate. Cells were treated with vehicle/ salinomycin (5 µM)/cisplatin (5 µM) and allowed to grow under 3D condition. After 72 hours, their corresponding photo micrographs are shown.(G) The effect of salinomycin and cisplatin in cell viability of spheroids were determined by trypan blue assay. Data points are average of triplicate readings of samples; error bars, ± S.D. *p < 0.01, compared to vehicle-treated cells. (H) Represents the FACS dot plot analysis of Annexin-V staining in CD133 gated population in the vehicle/salinomycin (10 µM)/cisplatin (10 µM) treated SW620 cells. Results shown from (A) to (H) sections are representative of at least three independent experiments.

Next, by utilizing this cellular model system with differential level of stemness, we sought to determine the *in-vitro* cytotoxic efficacy of both salinomycin and cisplatin in DLD-1 (CSC ^low^) and SW620 (CSC ^high^) cells by standard SRB assay. Interestingly, we observed that salinomycin posed robust cytotoxic effects against both the cell types whereas; cisplatin displayed cytotoxicity only against CSC ^low^ DLD-1 cells (Figure 1D). Transformation of anchorage-dependent (2D) growth to anchorage-independent (3D) growth or spheroid formation is a well-established characteristic feature of CSC development. Here, we first tested multiple colon cancer cells for spheroid formation and checked their stemness/differentiation status in 2D to 3D conditions. CSC ^low^ DLD-1 cells were found to be most suitable for our purpose as it showed reverse regulation in stemness (CD133) marker versus colonic epithelial differentiation (cytokeratin 20 or CK20) during 2D-3D transformation as observed by Western blot analysis (Figure 1E). So, we sought to determine the effect of both salinomycin and cisplatin in 2D-3D transformation of DLD-1 cells.

Interestingly, we observed that salinomycin effectively attenuated the spheroid formation, whereas, cisplatin treatment resulted in even larger spheroids instead of its inhibition (Figure 1F). We checked the viability control and treated cells, isolated from colonies and confirmed that salinomycin was able to induce cell death in spheroid colonies, whereas, cisplatin remained ineffective in killing CSCs (Figure 1G). So, these results together suggest that unlike chemotherapeutic drug cisplatin, salinomycincan effectively induces cytotoxicity to colon CSCs. Differential cytotoxic effect of salinomycin and cisplatin on colon CSCs prompted us to investigate whether these two can induce apoptosis in colon CSCs. To study the effects of salinomycin and cisplatin specifically on CSCs, we double stained our treated and control cells with CD133 (as stemness marker) and Annexin-V (as early apoptotic marker) and analyzed by flow cytometery to study the event of early apoptosis in stem cell (CD133^+^) compartment.

Interestingly, in the case of salinomycin treatment, we found a marked amount of Annexin-V^+^ early apoptotic cells in CD133^+^, but cisplatin failed to induce early apoptosis (Annexin-V^+^) in the same population (Figure 1H). Altogether, these results indicate that unlike the conventional chemotherapeutic drug cisplatin, salinomycin inhibits CSC properties and promotes apoptosis in colon CSCs.

### 3.2 Salinomycin selectively up-regulates the expression of functional DR4 and DR5 in colon CSCs and sensitizes them for TRAIL-induced apoptosis

After confirming that salinomycin induces apoptosis in colon CSCs (4, 17), we next explored the involvement of specific pathway for salinomycin induced apoptosis by using human apoptotic proteome profiler array (18). Array results in control versus salinomycin treatment indicate a change in the level of multiple proteins (Figure 2A-2B). The table shown in Figure 2A represents the identity of individual spots on the protein array platform. Based on densitometry of corresponding spots, we prepared a heat map to show relative changes in the level of corresponding protein (Figure 2B). Here, we found a marked increase in the level of DR4 and DR5 proteins in response to salinomycin treatment whereas; there was no change in expression of other signatory proteins like cytochrome C, Bax, and Bcl2 (Figure 2A-2B). Our array results were validated by performing individual Western blot analysis for the expression of DR4 and DR5 in both SW620 and LOVO cells (Figure 2C and Supplementary Figure 3), where robust up-regulation of both of these proteins were observed in two different doses (5 and 10 μM) of salinomycin treatment compared to vehicle-treated cells. To determine the salinomycin mediated DR4/DR5 up-regulation in colon CSCs (CD133^+^ cells), we performed dual staining of DR4-CD133 and DR5-CD133 in control and treated cells and analyzed by flow cytometry. As shown in histogram overlays, salinomycin treatment in both SW620 and LOVO cells resulted in strong upregulation of DR4 and DR5 on the cell surface of gated CD133^+^ population (Figure 2D). To validate the functional importance of this up-regulation of DR4/DR5 on the cell surface, we have analyzed the effect of human recombinant tumor necrosis factor (TNF)-related apoptosis-inducing ligand (TRAIL; ligand of DR4 and DR5) after 12 hours pre-treatment of salinomycin in both SW620 and LOVO cells which are classically known TRAIL-resistant cells. Here, we found that compared to respective controls, a short exposure of minimal doses (1, 2.5 μM) of salinomycin drastically increased the sensitivity of these cells towards TRAIL-mediated apoptosis (Figure 2E). Together, these data confirmed that salinomycin increases functional protein expression level of DR4 and DR5 on the cell surface of the colon CSCs and able to induce the apoptosis in the presence of TRAIL.

**Figure 2.**
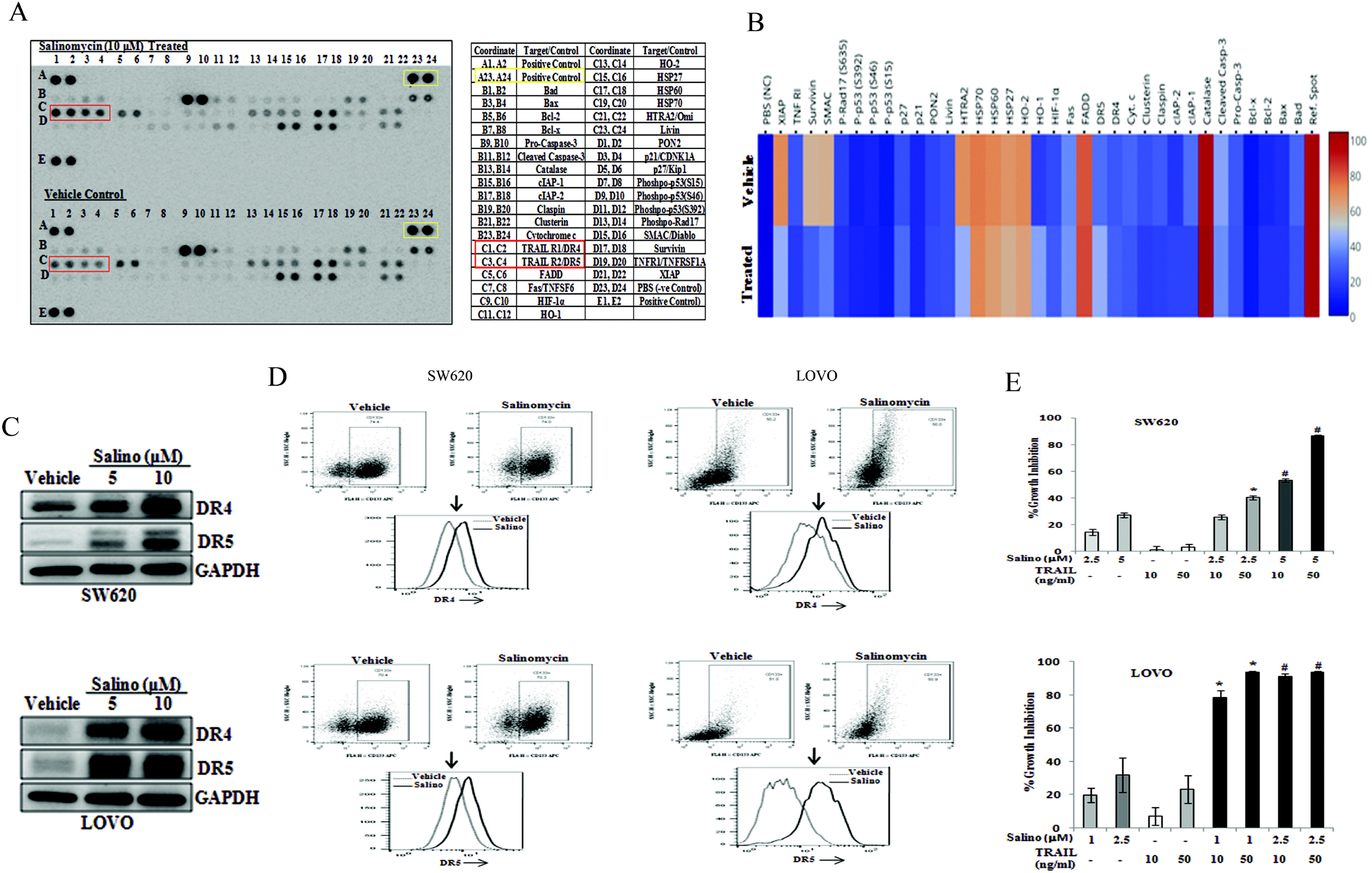
Salinomycin promotes the expression of DR4 and DR5 in colon cancer stem cells and sensitizes them for TRAIL-induced apoptosis. (A) Human proteome profiler apoptosis array was performed in vehicle and 10µM of salinomycin treated SW620 cells. Array spot coordinates for target proteins in duplicates were enlisted in right-hand panel. (B) Shows the heat map of respective proteins based on the pixel density of corresponding dots in vehicle and salinomycin treated groups. (C) The Western blot analysis of DR4, DR5 and GAPDH in vehicle and salinomycin treated SW620 (top) LOVO (bottom) cells; representative of at least three independent experiments. (D) SW620 and LOVO cells were treated with vehicle and salinomycin (10µM) for 24 hours and dual stained with either fluorochrome-conjugated CD133 or DR4 or CD133 and DR5 antibodies or matched isotype control and subjected to FACS analysis. Histogram overlays displaying the expression of either DR4 (top) or DR5 (bottom) in CD133 gated population of SW620 and LOVO cells, representative of at least three independent experiments. (E) SW620 (top) and LOVO (bottom) were pre-treated with salinomycin for 12 hours followed by TRAIL treatment for 36 hours and subjected to SRB assay. Results are representative of three independent experiments. Columns, an average of triplicate readings of samples; error bars, ± S.D. *, p < 0.05, compared with only 10ng/ml TRAIL treated cells; whereas, #, p < 0.05, compared with only 50ng/ml TRAIL treated cells.

### 3.3 Salinomycin up-regulates the expression of DR4 and DR5 by targeting epigenetic modulator EZH2

To investigate the molecular mechanism of salinomycin driven DR4 and DR5 up-regulation; we first, determined the effect of salinomycin in DR4 and DR5 transcription. So, we performed the RT-PCR in-vehicle control, and salinomycin treated SW620 cells. Our RT-PCR result suggests that salinomycin significantly up-regulates the mRNA expression of both DR4 and DR5 as compared to control (Figure 3A). Next, we analyzed whether salinomycin mediated induction in the expression of DR4/DR5is mediated by targeting of transcriptional regulators of *DR4/DR5* genes. P53 and SP1are known to act as an activator, while YY1 acts as a repressor of *DR4* and *DR5* gene transcription^31–34^ Here, we determine the expression of P53, SP1 and YY1 in vehicle control and salinomycin treated SW620 cells using Western blot. Surprisingly, our Western blot analysis of control and salinomycin treated cells indicates that there was no change in the SP1 level whereas p53 level was decreased after salinomycin treatment (Figure 3B).

**Figure 3.**
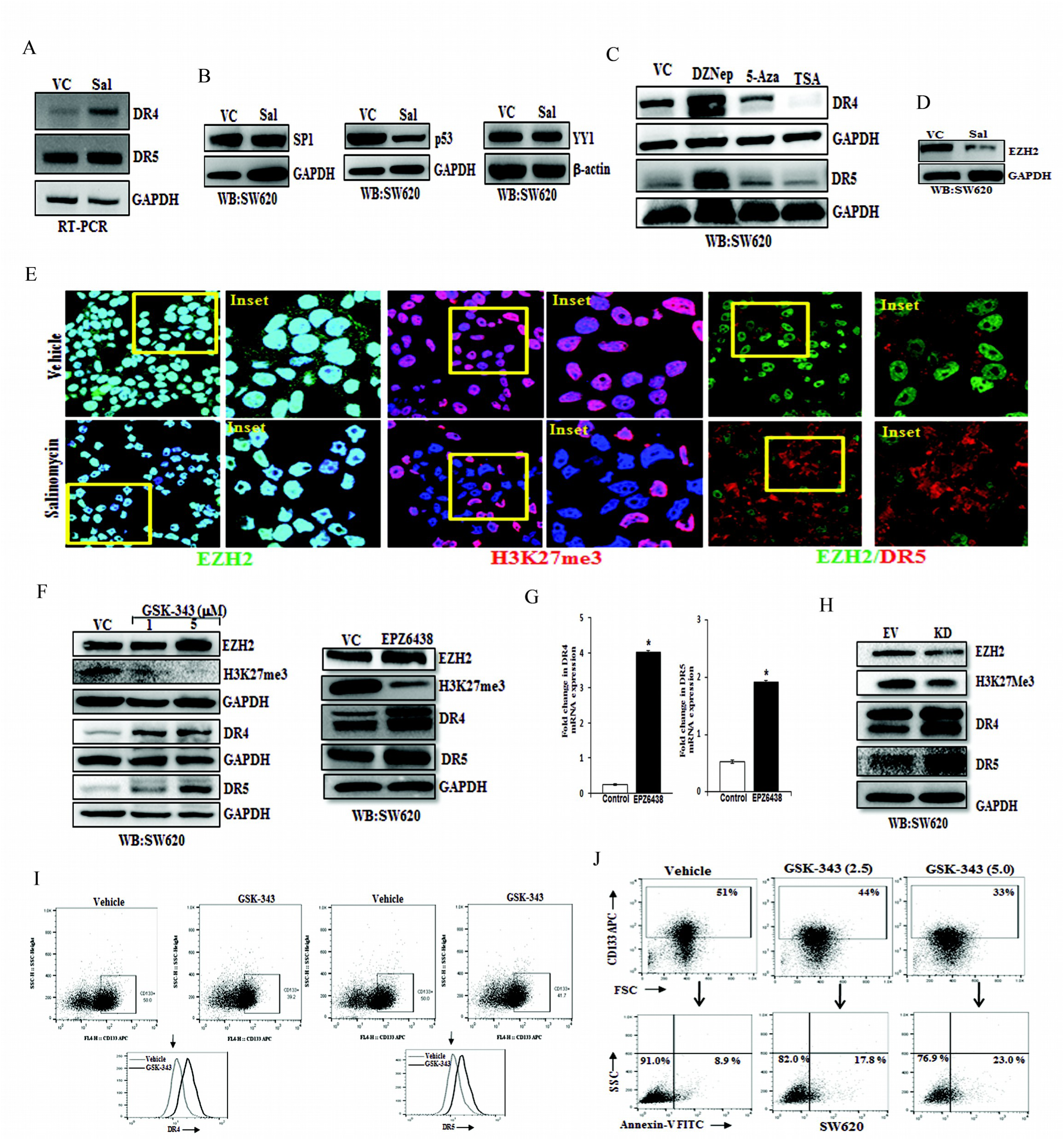
Salinomycin up-regulates the transcriptional expression of *DR4* and *DR5* genes by targeting EZH2. SW620 cells were treated with vehicle and salinomycin (10 μΜ) for 24 hours and subjected to (A) RT-PCR analysis. (B) Western blot for SP1, p53, YY1, GAPDH, β-actin. (C) SW620 cellswere treated with DzNep (10 μΜ), 5-Azacytidine (5 μΜ), Trichostatin-A (1 μΜ) for 24 hours followed by western blot analysis for DR4, DR5 and GAPDH protein expression. (D) SW620 cells were treated with vehicle and salinomycin (10 μΜ) followed by western blotting for EZH2 and GAPDH. (E) Confocal microscopy was performed for corresponding changes in EZH2 and global H3K27me3 level after vehicle or 10 µM salinomycin treatment for 24 hours. In similar setting, simultaneous expression of EZH2 and DR5 were assessed by confocal microscopy (Extreme right panel). Results shown in A-E are representative of at least three independent experiments. (F) SW620 cells were treated with 1 and 5 µM doses of GSK-343 and 10 µM dose of EPZ6438 and examine the expression of EZH2, H3K27me3, DR4, DR5 and GAPDH by western blot. (G) SW620 cells were treated with either vehicle or 10 µM dose of EPZ6438 and performed q-PCR analysis for DR4 and DR5 expression. Data points are average of duplicate readings of samples; error bars, ± S.D. *pL 0.01, compared to vehicle treated cells.(H) Control (Empty Vector) and EZH2 knockdown stable cells (SW620) were analyzed for expression of EZH2, H3K27me3, DR4, DR5 and GAPDH by western blot. SW620 cells were treated with either vehicle or GSK-343 for 24 hours and (I) dual stained with either FITC conjugated Annexin-V, and APC conjugated CD133 or (J) PE-conjugated DR4/DR5 and APC conjugated CD133 antibodies followed by FACS analysis. Results shown from (A)-(J) are representative of at least three independent experiments.

Similarly, there was no significant change in the YY1 level. As we did not observe any significant difference in the expression of the above-mentioned transcription factors following salinomycin treatment, we focus on possible epigenetic regulators of death receptors, if any. Therefore, we analyzed whether salinomycin target any specific epigenetic modulator which might be involved in the regulation of DR4/ DR5 expression. In this direction, we first, examine the expression of DR4/DR5 in SW620 cells treated with different epigenetic inhibitors like HNMT inhibitor 3-DeazaneplanocinA (DZNep), DNMT inhibitor 5’-Azacytidine (5-AZA) and HDAC inhibitor Trichostatin A (TSA) using western blot.

Interestingly, the western blot data of control and salinomycin treated cells reveal that the selective inhibition of HNMT protein EZH2 but not DNMT or HDAC inhibitors resulted in up-regulation of both DR4 and DR5 proteins in SW620 cells (Figure 3C). As DZNep treatment selectively promotes induction of death receptors, we checked the effect of salinomycin on expression of EZH2 and detected significant reduction of EZH2 following salinomycin treatment as compared to control in SW620 and HT-29 cells (Figure 3D and Supplementary Figure 4). Next, we assessed the expression of EZH2 and its functional enzymatic product H3K27me3 in control and salinomycin treated SW620 cells using confocal microscopy. Here, we observed that salinomycin robustly decreased the level of EZH2 and H3K27me3 compared to control (Figure 3E, left and middle panel) indicating that it delineates severe impairment of EZH2 functionality. Next, to understand the direct correlation between EZH2 and DR5 at a single-cell level, we simultaneously stained control and treated SW620 cells with EZH2 and DR5, and analyzed their expression together by using confocal microscopy (Figure 3E, right panel). Here, we found that salinomycin treatment not only decreased the EZH2 level both in the cytosol and nucleus but also increased the DR5 expression in the same population of cells. These results indicate that salinomycin inhibits EZH2 function which may result in the induction of DR4/DR5 in colon CSCs.

### 3.4 Genetic and pharmacological inhibition of EZH2 results in up-regulation of DR4 and DR5 in the colon CSCs

Based on our confocal microscopy results, we hypothesized that EZH2 enzymatic activity regulates the expression of *DR4/DR5* genes. In our preliminary observations, we used DZNep as an EZH2 inhibitor but it is known to have off-target effects. To rule out the off-target effects of EZH2 pharmacological inhibitors, here we selected two potent EZH2 inhibitors, i.e. GSK343 and EPZ6438, having completely different structural pharmacophores to validate our biological observations associated with it^35^. We treated SW620 cells with the vehicle control and two doses (1 and 5µM) of GSK-343 for 24 hours and analyzed the expression of EZH2, H3K27me3, DR4, and DR5 by Western blot. As shown in Figure 3F (left and right panel), compared to vehicle-treated SW620 cells, GSK-343and EPZ6438 treatment reduced the expression of H3K27me3 and simultaneously markedly increased expression of both DR4 and DR5. Similar results were obtained by using EPZ6438 in HT-29 cells (Supplementary Figure 5). Further to check whether EPZ6438 mediated increase in DR4/DR5 was at the transcriptional level or not, we performed real-time q-PCR analysis in EPZ6438 treated SW620 cells. We found a substantial mRNA increase in both *DR4* and *DR5* genes (Figure 3G) following EZH2 inhibitor treatment. The q-PCR analysis also suggests that endogenous mRNA expression of *DR4* gene is significantly lower than *DR5* in control SW620 cells that could be the result of predominant epigenetic transcriptional suppression of the *DR4* gene over *DR5*. To confirm our pharmacological inhibitor-based observations, we utilized genetic knockdown approach to examine EZH2 dependent DR4 and DR5 regulation. Here, we made stable EZH2 knockdown SW620 cells by utilizing the retroviral transduction system, and then we analyzed the expression of EZH2, H3K27me3, DR4, and DR5 by Western blot. Compared to the control, EZH2 knockdown resulted in a marked decrease in H3K27me3 level with significant up-regulation of both DR4 and DR5 proteins (Figure 3H).

As the previous series of data were generated using whole-cell population of SW620, now we sought to determine the impact of EZH2 inhibitor in regulation of death receptors in CSC compartment. Here, we assessed the effect of GSK-343 on cell surface expression of DR4/DR5, particularly in colon CSCs. Similar to salinomycin, we also found functional inhibition of EZH2 by GSK-343 resulted in marked increase in the surface expression of DR4 and DR5 in CD133^+^ compared to control (Figure 3I; left and right panels). Next, we sought to specifically determine the effect of EZH2 inhibitor in inducing apoptosis of colon CSCs. Using previously established strategy, we again double-stained control and GSK-343 treated SW620 cells for CD133 and Annexin-V followed by Flowcytometry analysis to study the event of early apoptosis in CSC (CD133^+^) compartment. Interestingly, it was observed that like salinomycin, EZH2 inhibitor treatment caused a prominent increase in Annexin-V^+^ early apoptotic cells in CD133^+^ cells (Figure 3J). Overall, these results clearly indicate that EZH2 directly regulates the expression of DR4 and DR5 by epigenetic suppression of their expression in colon cancer stem cells.

### 3.4 By targeting EZH2, salinomycin withdraws H3K27me3 marks near the promoters of DR4 and DR5

In the previous experiments, we observed Salinomycin and EZH2 inhibitor EPZ6438 transcriptionally regulates the expression of DR4 and DR5.EZH2 is known to suppress its target gene by trimethylating H3K27 in the vicinity of their promoters^36^. Therefore, we performed the chromatin immunoprecipitation (ChIP) assay to delineate the H3K27me3 marks near *DR4* and *DR5* genes. We designed walking primers for *DR4* and *DR5* from the sliding window of 1500 bases upstream and downstream of transcription start site (TSS) by using the UCSC genome browser (https://genome.ucsc.edu/). We first examined the enrichment of H3K27me3 marks in *DR4* and *DR5* promoters using real time q-PCR. The real time q-PCR analysis of immune-precipitated DNA demonstrated a strong enrichment of H3K27me3 marks as compared to control immunoglobulin (IgG) particularly in the *DR4* primer walking experiment. Among all the primers, the highest enrichment of H3K27me3 marks for *DR4* were observed near the primer flanking region −27 bases from TSS followed by +522 bases (Figure 4A).

**Figure 4.**
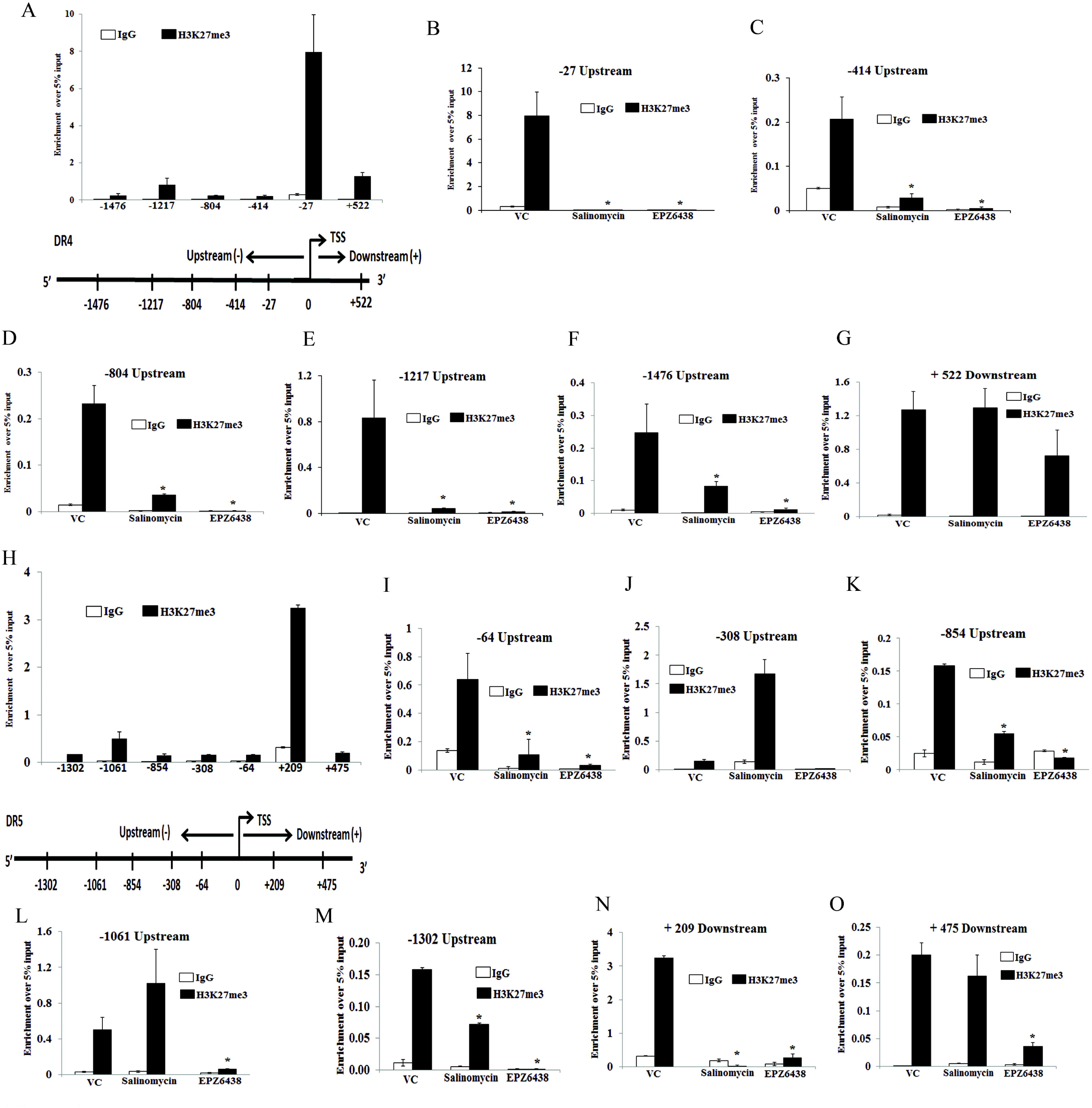
Salinomycin and EZH2 inhibitor remove H3K27me3 marks near the promoters of *DR4* and *DR5* genes. (A)H3K27me3 marks occupy DR4 promoter. ChIP was performed in SW620 cells using anti H3K27me3 and IgG antibodies and then examined by real-time q-PCR using primers pairs targeting −1.5 Kb to +1 Kb of the *DR4* gene. The X-axis indicates the central location of the PRC products relative to the DR4 TSS. (B-G) ChIP analysis showing differential fold change in H3K27me3 level at the promoter of *DR4* gene in SW620 cells treated with either vehicle or salinomycin (10 µM) or EPZ6438 (10µM) for 24 hours. (H) H3K27me3 marks occupy DR5 promoter. ChIP was performed in SW620 cells using anti H3K27me3 and IgG antibodies and then examined by real-time q-PCR using primers pairs targeting −1.5 Kb to +1 Kb of the *DR5* gene. The X-axis indicates the central location of the PRC products relative to the DR5 TSS. (I-O) ChIP analysis showing differential fold change in H3K27me3 level at the promoter of *DR5* in SW620 cells treated with either vehicle or salinomycin (10 µM) or EPZ6438 (10µM) for 24 hours. Results shown (A)-(O) are representative of two independent experiments. Columns, an average of duplicate readings of samples; error bars, ± S.D. *, p < 0.05, compared to vehicle treated cells.

Further, we separately treated SW620 cells with 5 μM of salinomycin and 10 μMofEPZ6438 and performed the ChIP assay and quantitatively analyzed the results by q-PCR. Our real-time q-PCR results show that both salinomycin and EPZ6438 were able to strongly eliminate the enrichment H3K27me3 marks from -27, -414, -804, and -1217 sites as compared to control (Figure 4B-4G). Of note, at certain sites like -1476 and +522, the EPZ6438 was found to be more potent to eliminate H3K27me3 marks. Subsequently, a similar experiment was performed for *DR5* to observe the distributions of H3K27me3 marks near its promoter. The ChIP-q-PCR results of *DR5* revealed that the major enrichment marks of H3K27me3 were present near the primer flanking region at position +209 as compared to the other sites (Figure 4H). Next, we analyzed the potential of both salinomycin and EPZ6438 to remove the H3k27me3 marks at DR5 promoter. Here, we found the elimination of H3K27me3 marks at a lesser number of sites that includes -854, -1302, and +475 (Figure 4I-4O). Altogether, our ChIP-qPCR results indicate that the enrichment marks of H3K27me3 near the DR4 promoter are higher as compared to DR5 in terms of the recruitment sites. Moreover, both salinomycin and EPZ6438 can significantly remove H3K27me3 repressive marks from the promoter regions of pro-apoptotic *DR4* and *DR5* genes.

### 3.6 EZH2 knockdown attenuates CSC properties *in-vitro* and *in-vivo*

To validate our *in*-*vitro* observations into *in*-*vivo*, we first developed DLD-1 xenograft model as it was observed to faithfully recapitulate functional CSC properties during 2D to 3D transformation. Utilizing the same model, we made multiple attempts to observe the *in-vivo* antitumor efficacy of salinomycin. Unfortunately, instead of efficacy, we found salinomycin treatment is extremely toxic to animals even at low dose as observed in earlier findings^37^. 2mg/kg intra-peritoneal daily dose of salinomycin resulted more than 20% weight loss in nude mice in a week time. Therefore, we decided to dissect the direct role of EZH2 in modulating CSC properties in colon cancer *in*-*vivo*, as EZH2 is the most viable drug target for salinomycin. Before going to *in-vivo* experiments, first we explored the impact of loss of EZH2 function in spheroid formation. So, we performed retroviral transduction to knock down EZH2 in DLD-1 cell line and performed Western blot analysis to confirm knockdown efficiency (Figure-5A). Next, we allowed both control vector and EZH2 knockdown cells to grow in the 3D culture condition. We observed that EZH2 knockdown markedly attenuates the spheroid formation, whereas, control cells formed the ideal spheroid colonies (Figure-5B). The trypan blue staining showed that the cell viability in EZH2 knockdown spheroid was significantly lower than that of control cell-derived spheroids (Figure-5C). The tumorigenic potential of cancer cell is another hallmark feature of CSC. To understand the influence of EZH2 in modulating *in-vivo* tumorigenic potential, we inoculated control and EZH2 knockdown DLD-1 cells at two different dilutions (0.5×10^6^ and 1×10^6^) in the right and left flank of same mice (n=3), respectively.

Interestingly, at lower dilution (0.5×10^6^), EZH2 knockdown DLD-1 cells failed to develop tumors in each of the three cases, whereas, the same number of control DLD1 inoculation resulted in prominent tumors in all three flanks suggesting strong inhibition of tumorigenic potential of colon cancer cells under EZH2 knockdown condition. In case of other dilution, EZH2 KD cell-derived tumors are smaller than control cell-derived tumors as demonstrated in representative pictures (Figure-5D). As observed in Figure 5E and 5F, inoculation of 2 million EZH2 KD cells resulted in significant (p<.01) reduction in tumor volume and weight as compared to control cell insulated tumors. Together, our results demonstrate that EZH2 plays a critical role in modulating CSC properties *in vitro* and *in vivo* in colon cancer.

**Figure 5.**
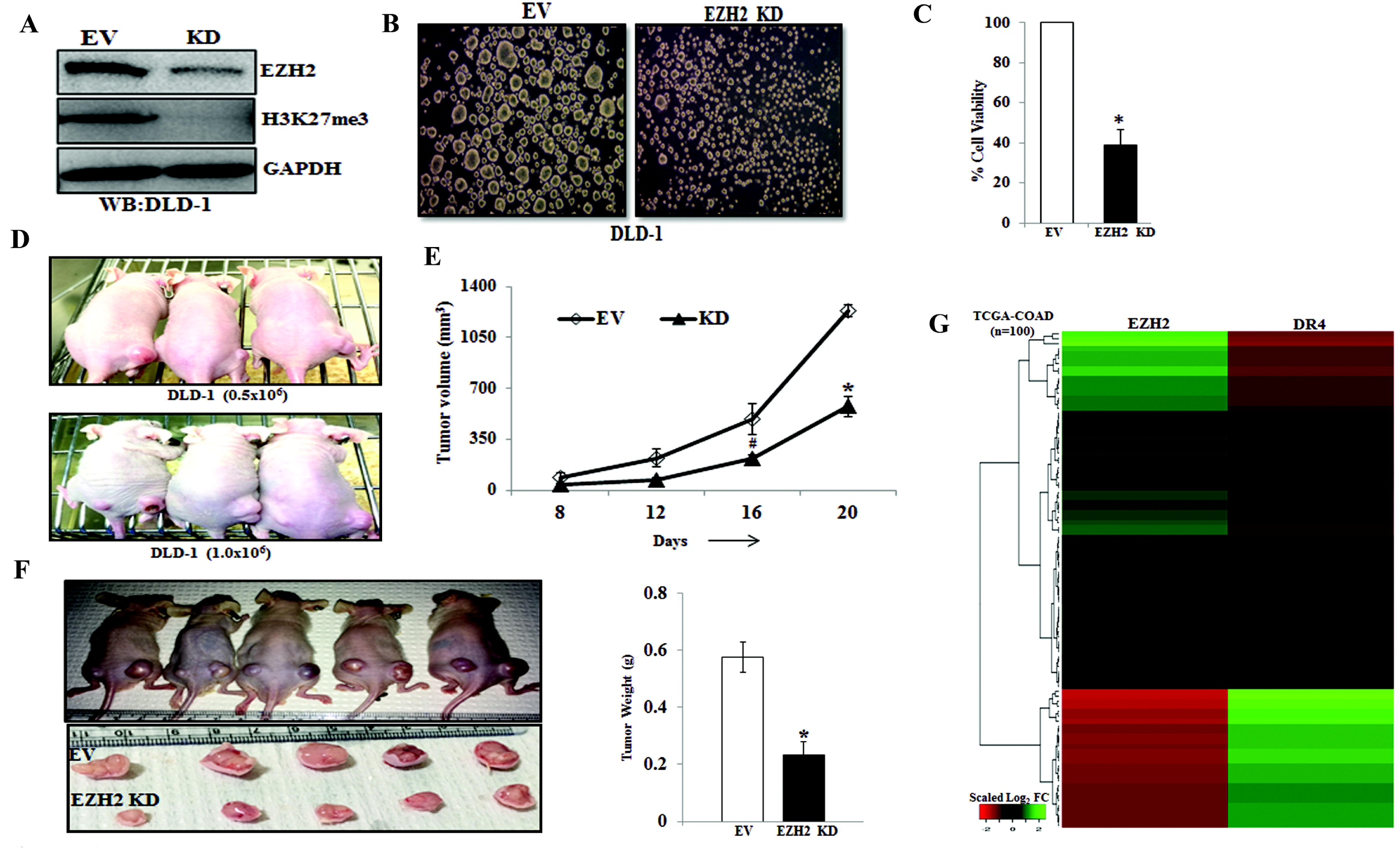
EZH2 knockdown inhibits CSC properties *in-vitro* and *in-vivo* and inversely correlated with DR4 expression. (A) Stable Empty Vector (EV) and EZH2 knockdown (KD) DLD-1 cells were analyzed for expression of EZH2, H3K27me3 and GAPDH by western blot. (B) EV and EZH2 KD DLD-1 cells were plated at 50,000 cells per 9.04 cm^2^ dish in a non-adherent plate and allowed to grow under 3D condition. After 72 hours, their corresponding photo micrographs are shown. (C) Trypan blue assay for percent cell viability in empty vector (EV) and EZH2 KD spheroid colonies. Data points are average of triplicate readings of samples; error bars, ± S.D. *pL 0.01, compared to control cells. Results shown from (A) to (C) sections are representative of at least three independent experiments. (D) EV and EZH2 KD DLD-1 cells at different dilutions (0.5×10^6^ and 1×10^6^) in 100μL PBS were subcutaneously inoculated in the right (EV) and left (KD) flank of each 4- to 6-week-old nude Crl: CD1-Foxn1^nu^ mice (n=3). After 2 weeks, photograph of tumor bearing mice were captured and represented. (E) EV and EZH2 KD DLD-1 cells (2×10^6^) in 100μL PBS were subcutaneously inoculated in the right (EV) and left (KD) flank of each 4- to 6-week-old nude Crl: CD1-Foxn1^nu^ mice (n=5) and allowed them grow for 21 days. Growth curve is shown for EV and EZH2 KD; points are indicative of average value of tumor volume; bars, +/- SD. The EZH2 KD group had significantly lower average tumor volumes from the EV group (^#^, p< 0.05; *, p< 0.01). (F) Representative images of tumor bearing mice and harvested tumors were shown (left panel). Average weight of harvested tumors bars, +/- SD of EV and EZH2 groups (*, p< 0.05) depicted by bar graph (right panel). (G) The heat map was generated to evaluate the correlation between EZH2 and DR4 mRNA signature in a segregated subset of high EZH2 (n=73) and low EZH2 (n=27) from COAD cohort of colon cancer patient data derived from TCGA database.

To understand the pathophysiological significance of our finding in context of human colon cancer, we exploited TCGA database to find out clinical correlation between EZH2 and death receptors. Interestingly, we observed a strong inverse correlation between EZH2 and DR4 expression in most of the colon cancer patient samples (Figure 5G), whereas, such inverse correlation is missing in case of DR5 expression (data not shown) again supporting our previous observations that EZH2 mediated DR4 regulation is predominant than DR5. Together, our results demonstrate that EZH2 can be therapeutically targeted to reduce CSC properties and DR4 expression may be a critical read out for inhibition of EZH2 function in human colon cancer.

## 4. Discussion

Though salinomycin has never positioned as a drug for cancer therapy, its discovery in the context of CSC targeting agent ignited new possibilities for finding novel molecular cues to tame therapy-resistant cancer cells. However, most of the studies for CSC specific cytotoxic function were based on genetically manipulated cells to develop a phenotype like CSCs in multiple cancers^7, 38^. For example, Gupta et al. enriched CSCs by loss of E-Cadherin function and tested the specificity of salinomycin to kill CSCs. In the course of dissecting salinomycin’s CSC specific cytotoxic function in physiologically relevant settings, we observed that salinomycin targets both CSC and non-CSC population which is corroborative for its immense toxic nature^37^. Nevertheless, the unique feature of salinomycin is that unlike standard chemotherapeutic drugs such as cisplatin, it can pose robust cytotoxic effects to cancer stem like cells.

We utilized this unique property of salinomycin to elucidate the mechanistic insight to target cancer stem cells or drug refractory cells. Earlier studies elucidated the influential role of high ALDH activity in drug resistance in general and high CD133 expression is associated with stemness in colon cancer^39, 40^. In alignment with previous results, we also observed the same reflection that salinomycin inhibits the expression of CD133 as well as ALDH activity in colon cancer cells. The flow-cytometry data demonstrated that salinomycin not only has the potential to induce apoptosis in CD133 enriched cells but also reduces CD133 expression to convert high CD133 enriched cells into low CD133enriched cells. Indeed, salinomycin can differentiate the CSCs to non-CSCs and supports the basic concept of CSC plasticity^25, 30, 40, 41^.

Recent studies suggest that the phenotypic changes through salinomycin treatment have linked with various signal transduction pathways like Wnt, K-RAS and Hedgehog signaling^42–44^. In addition, salinomycin targets various cellular processes in cancer which includes autophagy induction, mitochondrial impaired function, and depletion in ATP production. Besides, salinomycin treatment increases the production of reactive oxygen species (ROS) together with sequestration of iron in lysosomes, which induce ferroptosis in breast CSCs^45–49^.Functionally, salinomycin is a K^+^ selective ionophore; and it has been shown to act as passive potassium– hydrogen exchanger. Thus, facilitates the release of Ca^2+^ from ER to the cytosol and induce ER stress in cancer-like stem cells^50–53^. For the first time, here we put forward evidence for epigenetic basis of salinomycin CSC specific cytotoxic function. Our, unbiased human proteome profiler array data suggest that salinomycin targets *DR4/DR5* (key genes of extrinsic apoptotic pathways) and sensitize colon cancer stem cells for TRAIL therapy. Though the correlation between *DR4/DR5* with CSCs is dynamic and context-dependent, other findings suggest that CD133^+^ CSCs derived from human colon patients were resistant to TRAIL therapy^54^.

Several studies have demonstrated that salinomycin had synergistic effect with TRAIL therapy by DR5 modulation in glioma and ovarian cancers^55, 56^. Earlier finding regarding PRC2 mediated DR5 regulation is actually supporting our current observations^57^. However, histone methylation mediated epigenetic regulation of DR4 and the ability of salinomycin for TRAIL sensitization in colon CSCs was not reported so far. Though very limited information is present on the epigenetic regulation of *DR4/DR5*, prior studies examined that DR4 promoter was hyper-methylated in astrocytic glioma and DNMT inhibitor 5-aza-2-deoxycytidine treatment rescued the expression of *DR4* in a total population of different glioma cell lines^58^. However, in colon cancer cells, we did not find any correlation between promoter DNA methylation and DR4 expression. In our case, the treatment with 5-aza-2-deoxycytidine did not affect the expression of DR4/DR5 (Figure 3C). In fact, we analyzed the effect of direct modulation of all three epigenetic modifications including DNA methylation, histone methylation and histone acetylation on DR4/DR5 expression in an unbiased manner. This led us to discover that only histone methyl transferase EZH2 regulates the DR4/DR5 expression at transcriptional level. Thus, epigenetic landscapes are highly dynamic and context-dependent. The several reports suggested that EZH2 suppressed the expression of genes by tri-methylation of histone H3 at lysine 27 residue near the promoter of the target genes^19^. In support with our findings, Yang *et.al*., showed HOTAIR (non-coding RNA) regulates the expression of DR5 in pancreatic cancer *via* EZH2^59^.

Our extensive ChIP analysis demonstrated that H3K27me3 marks are highly enriched near the promoters of both *DR4* and *DR5* genes. Surprisingly, the salinomycin was not able to eliminate the H3K27me3 marks from all H3K27me3 enriched sites compared to the EZH2 inhibitor EPZ6438 in *DR4/DR5* genes. Therefore, EPZ6438, currently in Phase-II clinical trial, could be a wonderful therapeutics option to target CSCs instead of toxic salinomycin. ChIP data further confirmed that the epigenetic control by H3K27me3 is more predominant for DR4 as compared to DR5. Subsequently, this regulation was validated in colon CSCs, where functional diminution of EZH2 through pharmacological inhibition resulted in CD133 down-regulation as well as DR4/DR5 up-regulation in CD133^+^ cells. Consistent with our data; few recent studies have linked EZH2 with CSCs maintenance, activation and drug-resistant properties^60^. Previous studies described that EZH2 facilitates the expansion of breast stem cells through activation of NOTCH1 signaling and also maintained pancreatic cancer stem cells^61^.

Interestingly, it has been shown that EZH2 knockdown resulted in loss of drug-resistant side population or SP (CSC marker) and other CSC properties in ovarian cancer^62^. Another study showed that EZH2 is required for the stem cells regulation and tumorigenesis in skin cancer^63^.

In current study, EZH2 knock-down attenuated spheroid formation and suppressed the tumorigenic potential of colon cells in mouse xenograft model. Altogether, these findings delineate the central role of EZH2 in regulation of CSC phenotype, drug resistance and tumorigenesis and indeed, strongly advocating the immense potential of EZH2 inhibitors to overcome therapy resistance in cancer, especially by targeting CSC population. As a matter of fact, several EZH2 inhibitors like EPZ6438 and GSK126 are rapidly moving forward to Phase-I and II clinical trials against lymphoma and solid tumors^64^. In summary, our present findings provide mechanistic insights for the drug resistance properties in colon CSCs and suggest possible therapeutic interventions to overcome it. Nonetheless, it is the first report regarding the link between EZH2 and DR4 in colon cancer stem cells.

## 5. Acknowledgments

We sincerely acknowledge the excellent technical help of Mr. Sanjeev Meena for providing the routine cell culture facilities. Research of all the authors’ laboratories was supported by CSIR-CDRI Institutional Fund and Fellowship grants from CSIR, DBT and UGC. D.D acknowledges grant support from DST (EMR/2016/006935) and DBT (BT/AIR0568/PACE-15/18). Institutional (CSIR-CDRI) communication number for this article is 155.

## 7. Author Contributions

AKS and AV involved in study designing, performed experiments and wrote the draft manuscript. AS, SM, RKA, PC, MAN, KKS, JS provided active support for carrying out various *in-vitro* and *in-vivo* experiments. ALV helped in acquiring FACS data whereas, KS assisted for capturing confocal images. DD conceived the idea, designed experiments, analyzed data, wrote the manuscript and provided overall supervision. All authors read and approved the final manuscript.

## Data Availability

Data will be made available upon reasonable request.

## References

1. Bray, F., et al. Global cancer statistics 2018: GLOBOCAN estimates of incidence and mortality worldwide for 36 cancers in 185 countries. CA Cancer J. Clin. 68, 394–424, doi:10.3322/caac.21492 (2018).

2. De Angelis, M. L., Francescangeli, F., La Torre, F. & Zeuner, A. Stem Cell Plasticity and Dormancy in the Development of Cancer Therapy Resistance. Front. Oncol. 9, 626, doi:10.3389/fonc.2019.00626 (2019).

3. Martins-Neves, S. R., Cleton-Jansen, A. M. & Gomes, C. M. F. Therapy-induced enrichment of cancer stem-like cells in solid human tumors: Where do we stand? Pharmacol. Res. 137, 193–204, doi:10.1016/j.phrs.2018.10.011 (2018).

4. Ganesher, A. et al. New Spisulosine Derivative promotes robust autophagic response to cancer cells. Eur. J. Med. Chem. 188, 112011, doi:10.1016/j.ejmech.2019.112011 (2020).

5. Tiwari, R. et al. Androgen deprivation upregulates SPINK1 expression and potentiates cellular plasticity in prostate cancer. Nat Commun 11, 384, doi:10.1038/s41467-019-14184-0 (2020).

6. Singh, A. K. et al. Tumor heterogeneity and cancer stem cell paradigm: updates in concept, controversies and clinical relevance. Int. J. Cancer 136, 1991–2000, doi:10.1002/ijc.28804 (2015).

7. Gupta, P. B. et al. Identification of selective inhibitors of cancer stem cells by high-throughput screening. Cell 138, 645–659, doi:10.1016/j.cell.2009.06.034 (2009).

8. Lu, D. et al. Salinomycin inhibits Wnt signaling and selectively induces apoptosis in chronic lymphocytic leukemia cells. Proc. Natl. Acad. Sci. U. S. A. 108, 13253–13257, doi:10.1073/pnas.1110431108 (2011).

9. Easwaran, H., Tsai, H. C. & Baylin, S. B. Cancer epigenetics: tumor heterogeneity, plasticity of stem-like states, and drug resistance. Mol. Cell 54, 716–727, doi:10.1016/j.molcel.2014.05.015 (2014).

10. Munoz, P., Iliou, M. S. & Esteller, M. Epigenetic alterations involved in cancer stem cell reprogramming. Mol. Oncol. 6, 620–636, doi:10.1016/j.molonc.2012.10.006 (2012).

11. Widschwendter, M. et al. Epigenetic stem cell signature in cancer. Nat. Genet. 39, 157–158, doi:10.1038/ng1941 (2007).

12. Valk-Lingbeek, M. E., Bruggeman, S. W. & van Lohuizen, M. Stem cells and cancer; the polycomb connection. Cell 118, 409–418, doi:10.1016/j.cell.2004.08.005 (2004).

13. Francis, N. J., Kingston, R. E. & Woodcock, C. L. Chromatin compaction by a polycomb group protein complex. Science 306, 1574–1577, doi:10.1126/science.1100576 (2004).

14. Bhatia, V. et al. Epigenetic Silencing of miRNA-338-5p and miRNA-421 Drives SPINK1-Positive Prostate Cancer. Clin. Cancer Res. 25, 2755–2768, doi:10.1158/1078-0432.CCR-18-3230 (2019).

15. Cao, R. et al. Role of histone H3 lysine 27 methylation in Polycomb-group silencing. Science 298, 1039–1043, doi:10.1126/science.1076997 (2002).

16. Varambally, S. et al. The polycomb group protein EZH2 is involved in progression of prostate cancer. Nature 419, 624–629, doi:10.1038/nature01075 (2002).

17. Kleer, C. G. et al. EZH2 is a marker of aggressive breast cancer and promotes neoplastic transformation of breast epithelial cells. Proc. Natl. Acad. Sci. U. S. A. 100, 11606–11611, doi:10.1073/pnas.1933744100 (2003).

18. Suva, M. L. et al. EZH2 is essential for glioblastoma cancer stem cell maintenance. Cancer Res. 69, 9211–9218, doi:10.1158/0008-5472.CAN-09-1622 (2009).

19. Deb, G., Singh, A. K. & Gupta, S. EZH2: not EZHY (easy) to deal. Mol. Cancer Res. 12, 639–653, doi:10.1158/1541-7786.MCR-13-0546 (2014).

20. Khanbolooki, S. et al. Nuclear factor-kappaB maintains TRAIL resistance in human pancreatic cancer cells. Mol. Cancer Ther. 5, 2251–2260, doi:10.1158/1535-7163.MCT-06-0075 (2006).

21. Zhang, Y. & Zhang, B. TRAIL resistance of breast cancer cells is associated with constitutive endocytosis of death receptors 4 and 5. Mol. Cancer Res. 6, 1861–1871, doi:10.1158/1541-7786.MCR-08-0313 (2008).

22. Singh, A. K. et al. Dual targeting of MDM2 with a novel small-molecule inhibitor overcomes TRAIL resistance in cancer. Carcinogenesis 37, 1027–1040, doi:10.1093/carcin/bgw088 (2016).

23. Fulda, S. & Debatin, K. M. Exploiting death receptor signaling pathways for tumor therapy. Biochim. Biophys. Acta 1705, 27–41, doi:10.1016/j.bbcan.2004.09.003 (2004).

24. Vichai, V. & Kirtikara, K. Sulforhodamine B colorimetric assay for cytotoxicity screening. Nat. Protoc. 1, 1112–1116, doi:10.1038/nprot.2006.179 (2006).

25. Arya, R. K. et al. Anti-breast tumor activity of Eclipta extract in-vitro and in-vivo: novel evidence of endoplasmic reticulum specific localization of Hsp60 during apoptosis. Sci. Rep. 5, 18457, doi:10.1038/srep18457 (2015).

26. Datta, D. et al. Ras-induced modulation of CXCL10 and its receptor splice variant CXCR3-B in MDA-MB-435 and MCF-7 cells: relevance for the development of human breast cancer. Cancer Res. 66, 9509–9518, doi:10.1158/0008-5472.CAN-05-4345 (2006).

27. Maheshwari, S. et al. Discovery of a Novel Small-Molecule Inhibitor that Targets PP2A-beta-Catenin Signaling and Restricts Tumor Growth and Metastasis. Mol. Cancer Ther. 16, 1791–1805, doi:10.1158/1535-7163.MCT-16-0584 (2017).

28. Dillies, M. A. et al. A comprehensive evaluation of normalization methods for Illumina high-throughput RNA sequencing data analysis. Brief. Bioinform. 14, 671–683, doi:10.1093/bib/bbs046 (2013).

29. Dewangan, J., Srivastava, S. & Rath, S. K. Salinomycin: A new paradigm in cancer therapy. Tumour Biol. 39, 1010428317695035, doi:10.1177/1010428317695035 (2017).

30. Chaffer, C. L. et al. Normal and neoplastic nonstem cells can spontaneously convert to a stem-like state. Proc. Natl. Acad. Sci. U. S. A. 108, 7950–7955, doi:10.1073/pnas.1102454108 (2011).

31. Liu, X., Yue, P., Khuri, F. R. & Sun, S. Y. p53 upregulates death receptor 4 expression through an intronic p53 binding site. Cancer Res. 64, 5078–5083, doi:10.1158/0008-5472.CAN-04-1195 (2004).

32. Takimoto, R. & El-Deiry, W. S. Wild-type p53 transactivates the KILLER/DR5 gene through an intronic sequence-specific DNA-binding site. Oncogene 19, 1735–1743, doi:10.1038/sj.onc.1203489 (2000).

33. Yoshida, T., Maeda, A., Tani, N. & Sakai, T. Promoter structure and transcription initiation sites of the human death receptor 5/TRAIL-R2 gene. FEBS Lett. 507, 381–385, doi:10.1016/s0014-5793(01)02947-7 (2001).

34. Martinez-Paniagua, M. A. et al. Mcl-1 and YY1 inhibition and induction of DR5 by the BH3-mimetic Obatoclax (GX15-070) contribute in the sensitization of B-NHL cells to TRAIL apoptosis. Cell Cycle 10, 2792–2805, doi:10.4161/cc.10.16.16952 (2011).

35. Knutson, S. K. et al. Selective inhibition of EZH2 by EPZ-6438 leads to potent antitumor activity in EZH2-mutant non-Hodgkin lymphoma. Mol. Cancer Ther. 13, 842–854, doi:10.1158/1535-7163.MCT-13-0773 (2014).

36. Yu, J. et al. The neuronal repellent SLIT2 is a target for repression by EZH2 in prostate cancer. Oncogene 29, 5370–5380, doi:10.1038/onc.2010.269 (2010).

37. Boehmerle, W., Muenzfeld, H., Springer, A., Huehnchen, P. & Endres, M. Specific targeting of neurotoxic side effects and pharmacological profile of the novel cancer stem cell drug salinomycin in mice. J. Mol. Med. (Berl.) 92, 889–900, doi:10.1007/s00109-014-1155-0 (2014).

38. Naujokat, C. & Steinhart, R. Salinomycin as a drug for targeting human cancer stem cells. J. Biomed. Biotechnol. 2012, 950658, doi:10.1155/2012/950658 (2012).

39. Zhi, Q. M. et al. Salinomycin can effectively kill ALDH(high) stem-like cells on gastric cancer. Biomed. Pharmacother. 65, 509–515, doi:10.1016/j.biopha.2011.06.006 (2011).

40. Gupta, P. B. et al. Stochastic state transitions give rise to phenotypic equilibrium in populations of cancer cells. Cell 146, 633–644, doi:10.1016/j.cell.2011.07.026 (2011).

41. Tang, D. G. Understanding cancer stem cell heterogeneity and plasticity. Cell Res. 22, 457–472, doi:10.1038/cr.2012.13 (2012).

42. Najumudeen, A. K. et al. Cancer stem cell drugs target K-ras signaling in a stemness context. Oncogene 35, 5248–5262, doi:10.1038/onc.2016.59 (2016).

43. Lu, Y. et al. Salinomycin exerts anticancer effects on human breast carcinoma MCF-7 cancer stem cells via modulation of Hedgehog signaling. Chem. Biol. Interact. 228, 100–107, doi:10.1016/j.cbi.2014.12.002 (2015).

44. Huang, X. et al. The Molecular Basis for Inhibition of Stemlike Cancer Cells by Salinomycin. ACS Cent Sci 4, 760–767, doi:10.1021/acscentsci.8b00257 (2018).

45. Jangamreddy, J. R. et al. Salinomycin induces activation of autophagy, mitophagy and affects mitochondrial polarity: differences between primary and cancer cells. Biochim. Biophys. Acta 1833, 2057–2069, doi:10.1016/j.bbamcr.2013.04.011 (2013).

46. Kim, S. H. et al. Salinomycin simultaneously induces apoptosis and autophagy through generation of reactive oxygen species in osteosarcoma U2OS cells. Biochem. Biophys. Res. Commun. 473, 607–613, doi:10.1016/j.bbrc.2016.03.132 (2016).

47. Boehmerle, W. & Endres, M. Salinomycin induces calpain and cytochrome c-mediated neuronal cell death. Cell Death Dis. 2, e168, doi:10.1038/cddis.2011.46 (2011).

48. Verdoodt, B. et al. Salinomycin induces autophagy in colon and breast cancer cells with concomitant generation of reactive oxygen species. PLoS One 7, e44132, doi:10.1371/journal.pone.0044132 (2012).

49. Mai, T. T. et al. Salinomycin kills cancer stem cells by sequestering iron in lysosomes. Nat. Chem. 9, 1025–1033, doi:10.1038/nchem.2778 (2017).

50. Heijmans, J. et al. ER stress causes rapid loss of intestinal epithelial stemness through activation of the unfolded protein response. Cell Rep. 3, 1128–1139, doi:10.1016/j.celrep.2013.02.031 (2013).

51. Wielenga, M. C. B. et al. ER-Stress-Induced Differentiation Sensitizes Colon Cancer Stem Cells to Chemotherapy. Cell Rep. 13, 489–494, doi:10.1016/j.celrep.2015.09.016 (2015).

52. Feng, Y. X. et al. Epithelial-to-mesenchymal transition activates PERK-eIF2alpha and sensitizes cells to endoplasmic reticulum stress. Cancer Discov. 4, 702–715, doi:10.1158/2159-8290.CD-13-0945 (2014).

53. Mekahli, D., Bultynck, G., Parys, J. B., De Smedt, H. & Missiaen, L. Endoplasmic-reticulum calcium depletion and disease. Cold Spring Harb. Perspect. Biol. 3, doi:10.1101/cshperspect.a004317 (2011).

54. Todaro, M. et al. Colon cancer stem cells dictate tumor growth and resist cell death by production of interleukin-4. Cell Stem Cell 1, 389–402, doi:10.1016/j.stem.2007.08.001 (2007).

55. Calzolari, A. et al. Salinomycin potentiates the cytotoxic effects of TRAIL on glioblastoma cell lines. PLoS One 9, e94438, doi:10.1371/journal.pone.0094438 (2014).

56. Parajuli, B. et al. Salinomycin induces apoptosis via death receptor-5 up-regulation in cisplatin-resistant ovarian cancer cells. Anticancer Res. 33, 1457–1462 (2013).

57. Benoit, Y. D., Laursen, K. B., Witherspoon, M. S., Lipkin, S. M. & Gudas, L. J. Inhibition of PRC2 histone methyltransferase activity increases TRAIL-mediated apoptosis sensitivity in human colon cancer cells. J. Cell. Physiol. 228, 764–772, doi:10.1002/jcp.24224 (2013).

58. Elias, A. et al. Epigenetic silencing of death receptor 4 mediates tumor necrosis factor-related apoptosis-inducing ligand resistance in gliomas. Clin. Cancer Res. 15, 5457–5465, doi:10.1158/1078-0432.CCR-09-1125 (2009).

59. Yang, S. Z. et al. The long non-coding RNA HOTAIR enhances pancreatic cancer resistance to TNF-related apoptosis-inducing ligand. J. Biol. Chem. 292, 10390–10397, doi:10.1074/jbc.M117.786830 (2017).

60. Hu, S. et al. Overexpression of EZH2 contributes to acquired cisplatin resistance in ovarian cancer cells in vitro and in vivo. Cancer Biol. Ther. 10, 788–795, doi:10.4161/cbt.10.8.12913 (2010).

61. van Vlerken, L. E. et al. EZH2 is required for breast and pancreatic cancer stem cell maintenance and can be used as a functional cancer stem cell reporter. Stem Cells Transl Med 2, 43–52, doi:10.5966/sctm.2012-0036 (2013).

62. Rizzo, S. et al. Ovarian cancer stem cell-like side populations are enriched following chemotherapy and overexpress EZH2. Mol. Cancer Ther. 10, 325–335, doi:10.1158/1535-7163.MCT-10-0788 (2011).

63. Adhikary, G. et al. Survival of skin cancer stem cells requires the Ezh2 polycomb group protein. Carcinogenesis 36, 800–810, doi:10.1093/carcin/bgv064 (2015).

64. Kim, K. H. & Roberts, C. W. Targeting EZH2 in cancer. Nat. Med. 22, 128–134, doi:10.1038/nm.4036 (2016).

