## Supplementary Data for "Salinomycin inhibits epigenetic modulator EZH2 to enhance Death Receptors in Colon Cancer Stem Cells"

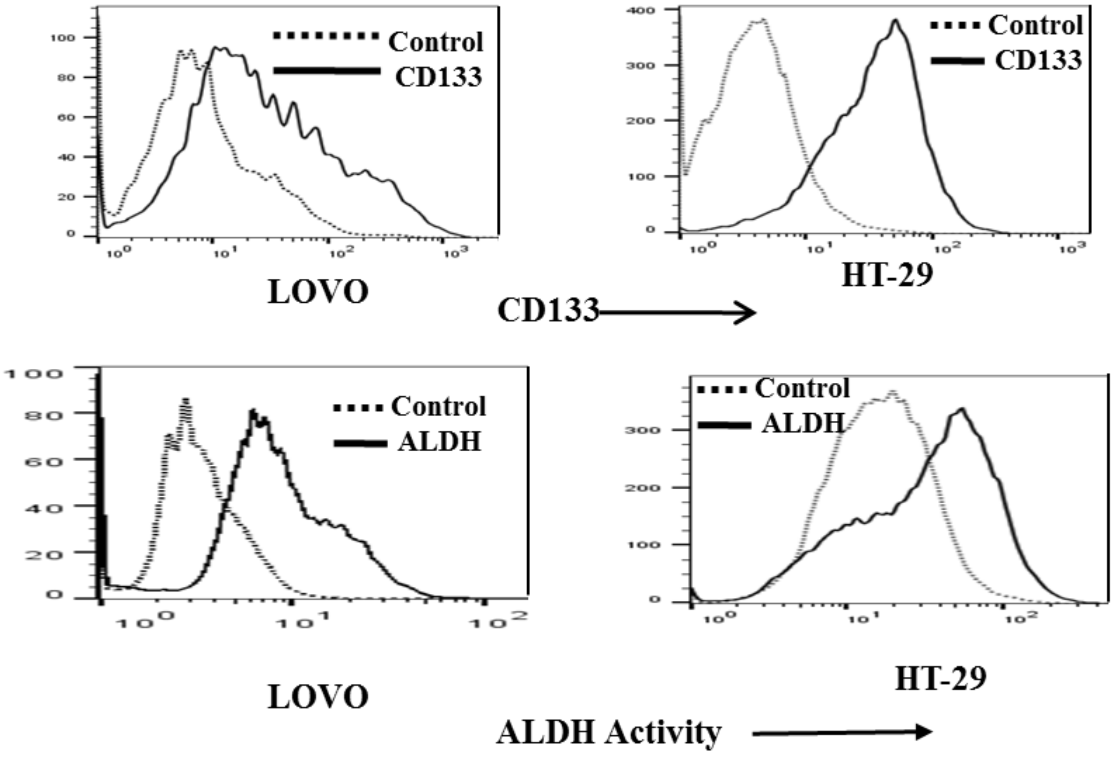


**Supplementary Figure- 1: CD133 surface expression and ALDH activity in LOVO and HT-29 cells**
Top Panel: CD133 surface expression were analyzed in LOVO, HT-29 cells by FACS
**Bottom Panel:** ALDH activity in LOVO, HT-29 cells was determined by FACS , as described Materials and Method section .

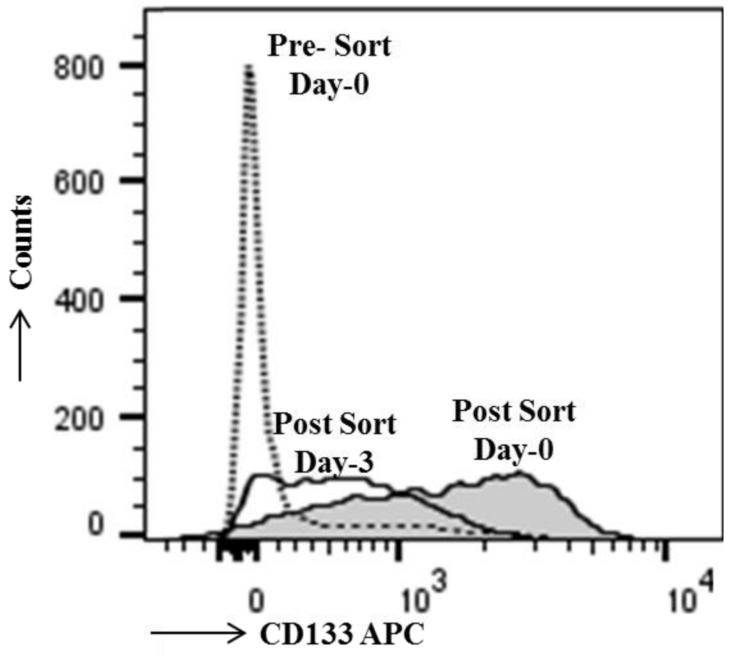


**Supplementary Figure- 2: CD133+ DLD-1 cells rapidly converted into CD133- cells and quickly lost their CSC integrity in cell culture.**

DLD-1 cells were stained with APC conjugated CD133 antibody, followed by FACS Aria based CD133+ cell sorting. Data were analyzed for presort and post sorted population. CD133+ sorted cells were cultured for 3 days and again stained with CD133 antibody and analyzed by flow-cytometry.

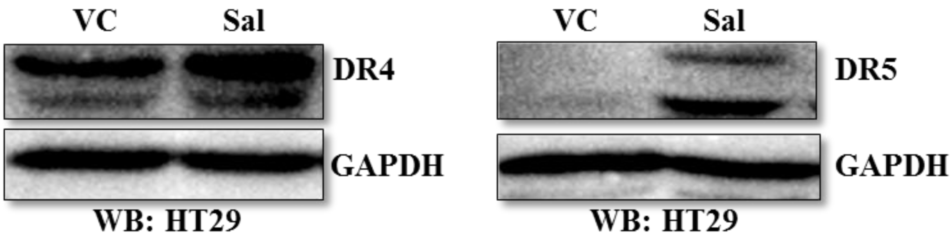


**Supplementary Figure-3: Salinomycin promotes expression of DR4 and DR5 in HT-29 cells**
HT-29 cells were treated with vehicle and salinomycin (10μm) dose for 24 hours. The western blot was performed for DR4 , DR5 and GAPDH. Results are representative of three independent experiments.

---

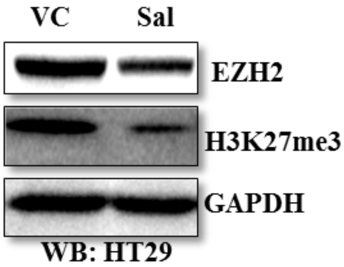


**Supplementary Figure-4: Salinomycin mediated down-regulation of EZH2 in HT-29 cells**
HT-29 cells were treated with vehicle and salinomycin (10μm) dose for 24 hours. The western blot was performed for EZH2, H3K27me3 and GAPDH. Results are representative of three independent experiments.

---

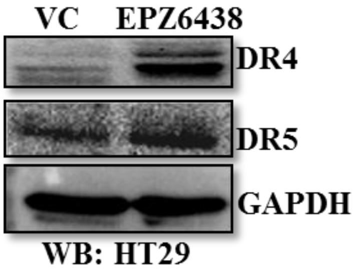


**Supplementary Figure-5: EZH2 inhibitor EPZ6438 treatment upregulates the expression of DR4 and DR5 in HT-29 cells**
HT-29 cells were treated with vehicle and 10 µM dose of EPZ6438 for 24 hours and examined the expression of DR4, DR5 and GAPDH by western blot. . Results are representative of three independent experiments.

**Supplementary Table- 1**

**ChIP primers list: DR4 and DR5**

| | |
|---|---|
| UA_ Forward _DR4 | CTACTCGGGAGGCTGAGGCA |
| UA_ Reverse_DR4 | AAACACACGGAGGCCAGCTC |
| UB_Forward_DR4 | TCCGACAGACTGGGGAGCAA |
| UB_Reverse_DR4 | TTCGGGAGGCCACTGGTGTA |
| UC_Forward _DR4 | ATGGTGGCCGTCCAGTAAGC |
| UC_ Reverse_DR4 | GATGAGAGCTGCCCACTGCC |
| UD_Forward_DR4 | AACACTTACGTGTGGCCGGG |
| UD_Reverse_DR4 | AGCTCAGCCTCCCAGGTTCA |
| UE_Forward_DR4 | ACTCCAGCCTGGGTGACAGA |
| UE_Reverse_DR4 | TCCTTGTGCATTTTGTTCTTCAGTTG |
| DA_Forward_DR4 | CCTCCCCAGGTGGGTTCTCA |
| DA_Reverse_DR4 | TCAGGCCCAGAACCCAAGGA |
| DB_Forward_DR4 | CGTGCCCTTTGACTCCTGGG |
| DB_Reverse_DR4 | GATCGAGGCGTTCCGTCCAG |
| UA_Forward_DR5 | ACAGGAAACCAGGGCTGTGC |
| UA_Reverse_DR5 | TGGTGTCCCTGTCCCCAGAG |
| UB_Forward_DR5 | CGGGTTGCATCTGGTTCCGA |
| UB_Reverse_DR5 | TGGGTACCCTGGTGGGGATG |
| UC_Forward_DR5 | AGCCGCCTGGTCTCTCTTGA |
| UC_Reverse_DR5 | GTTTTCCTGGAGCCCACCCC |
| UD_Forward_DR5 | CCTCCTCCGTGGGACTCCAT |
| UD_Reverse_DR5 | TGGGGGCTGGGAGCTAGTTT |
| UE_Forward_DR5 | GGCTGTGTCCTGTGGCCTTT |
| UE_Reverse_DR5 | AGGAAGGGAGCCTGCAGTCT |
| DA_Forward_DR5 | AGCGATTCTCGTGCCTCAGC |
| DA_Reverse_DR5 | TGATCCACCTTGGCCTCCGA |
| DB_Forward_DR5 | GCTTTGCTATATCCCCAGGCCA |
| DB_Reverse_DR5 | TCTGCCCTGTCAAAGGTCCCT |
